# Context-Dependent Adaptation in Structured Environments

**DOI:** 10.1101/2025.04.29.651346

**Authors:** Yuya Karita, Gisela T. Rodríguez-Sánchez, Elisa Brambilla, J. Carlos R. Hernandez-Beltran, Michael Schwarz, Paul B. Rainey

**Affiliations:** Department of Microbial Population Biology, Max Planck Institute for Evolutionary Biology, 24306 Plön, Germany; Department of Physics, The University of Tokyo, 113-0033 Tokyo, Japan; Laboratory of Biophysics and Evolution, CBI, ESPCI Paris, UniversitéPSL CNRS, 75005 Paris, France

## Abstract

*Adaptive evolution often leads to niche specialization, but successful colonization of a new niche can depend as much on ecological context as on genetic change. This is especially true in spatially structured environments, where the physical construction of habitat and the order of arrival of mutants can shape evolutionary outcomes. In static broth cultures, the air-liquid interface (ALI) provides a high-oxygen niche typically colonized by mat-forming mutants. In Pseudomonas fluorescens*, a range of single mutations that upregulate cellulose production give rise to “wrinkly spreader” (WS) types—canonical niche specialists that out-compete ancestral types at the ALI. Yet we discover that this is a half-truth: WS mutants frequently fail to establish when introduced alone at low density, but co-culture with ancestral types rescues colonization. Microscopy and model simulations show that ancestors physically scaffold the ALI, enabling stable attachment and facilitating the *in situ* emergence of an array of genetically diverse WS mutants. Ancestors later disperse, vacate the niche allowing WS mutant expansion. This transient asymmetric interaction reveals that even strongly adaptive mutations may require ecological facilitation to succeed. More broadly, our findings challenge simple narratives of adaptive replacement and underscore how transient ecological dependencies can shape the trajectory of evolutionary change.

## Introduction

The air–liquid interface (ALI) in static liquid cultures presents a distinct oxygen-rich niche that supports the adaptive evolution of various mat-forming bacteria^1–5^. In such spatially structured environments, aerobic growth rapidly depletes oxygen in the bulk liquid phase, creating steep vertical gradients that select for mutants capable of colonizing the ALI^6^. These mutants form substantive mats at the ALI, securing access to oxygen and a competitive edge in the population.

*Pseudomonas fluorescens* SBW25 has served as a model for studying this process. Under static conditions, the ancestral smooth morphotype (SM) cannot stably occupy the ALI, but de novo mutants known as “wrinkly spreaders” (WS) routinely evolve and dominate within days^1^. WS types over-produce extracellular matrices due to mutations that up-regulate cyclic dimeric guanosine monophosphate (c-di-GMP), a second messenger controlling the motile–sessile transition in many bacterial species^7–9^. These mutations have been mapped to multiple genetic pathways and occur at a combined rate of 1.1 *×* 10^−8^ [1/generation]^10^, enabling rapid and reproducible evolutionary change.

Yet despite this apparent advantage, WS types present several paradoxes. They grow more slowly than the ancestral SM type under both oxygen-rich and oxygen-depleted conditions^6,11^, and they exhibit reduced motility^12^. These traits should hinder, rather than promote, colonization of the ALI in inter-strain competition. Moreover, the ancestral SM type is itself capable of transient ALI colonization via the formation of thousands of microcolonies at the meniscus that over a matter of hours coalesce and collapse^13^. These observations challenge the assumption that WS mutants simply out-compete ancestral types due to superior niche adaptation.

Previous work has proposed that WS mutants succeed by providing collective benefits—specifically, oxygen access through mat formation—despite individual costs^11,14^. In this view, ancestral types in the mat are often treated as cheaters or free riders that exploit this structure without contributing to its formation^14,15^. While this population-level interpretation is supported by data ^11^, the microscopic dynamics of how slow-growing WS types emerge and spread remain poorly understood.

In this study, we revisit the dynamics of WS evolution with a focus on the ecological and spatial context in which WS mutants arise. We show that WS types, when introduced alone at low cell density, frequently fail to colonize the ALI within 24 hours. This deficiency is rescued by the presence of ancestral SM cells in co-culture, suggesting a previously unrecognized ecological interaction. Microscopy and modeling reveal that SM cells physically scaffold the ALI, compensating for WS motility defects and enabling stable attachment. Once established, WS types eventually displace ancestral types through differences in dispersal behavior—a “dispersal-driven” dynamic that contrasts with classical selective sweeps. These dynamics lead to the *in situ* emergence and coexistence of multiple WS lineages within a single population. Our results reveal a hidden dependency on ancestral types in what appears to be a simple case of adaptive radiation, and more broadly, highlight the role of ecological facilitation in shaping evolutionary outcomes in structured environments.

## Results

### Fitness costs associated with WS types

*P. fluorescens* SBW25 reproducibly generates mat-forming mutants in 3-6 days in static culture conditions (Fig. 1a). These wrinkly spreader (WS) mutants, termed based on their characteristic colony morphs^1^, are known to exhibit several fitness costs, including slower growth rates^6^ and reduced motility^12,16^. While impaired motility limits ability to reach the air–liquid interface (ALI), WS types are more hydrophobic than ancestral smooth colony morphotypes (SM)^17^, allowing them to raft more stably once at the surface. Thus, the fitness landscape for WS is shaped by a trade-off between impaired migration to, and enhanced persistence at, the ALI^12^.

**Figure 1.**
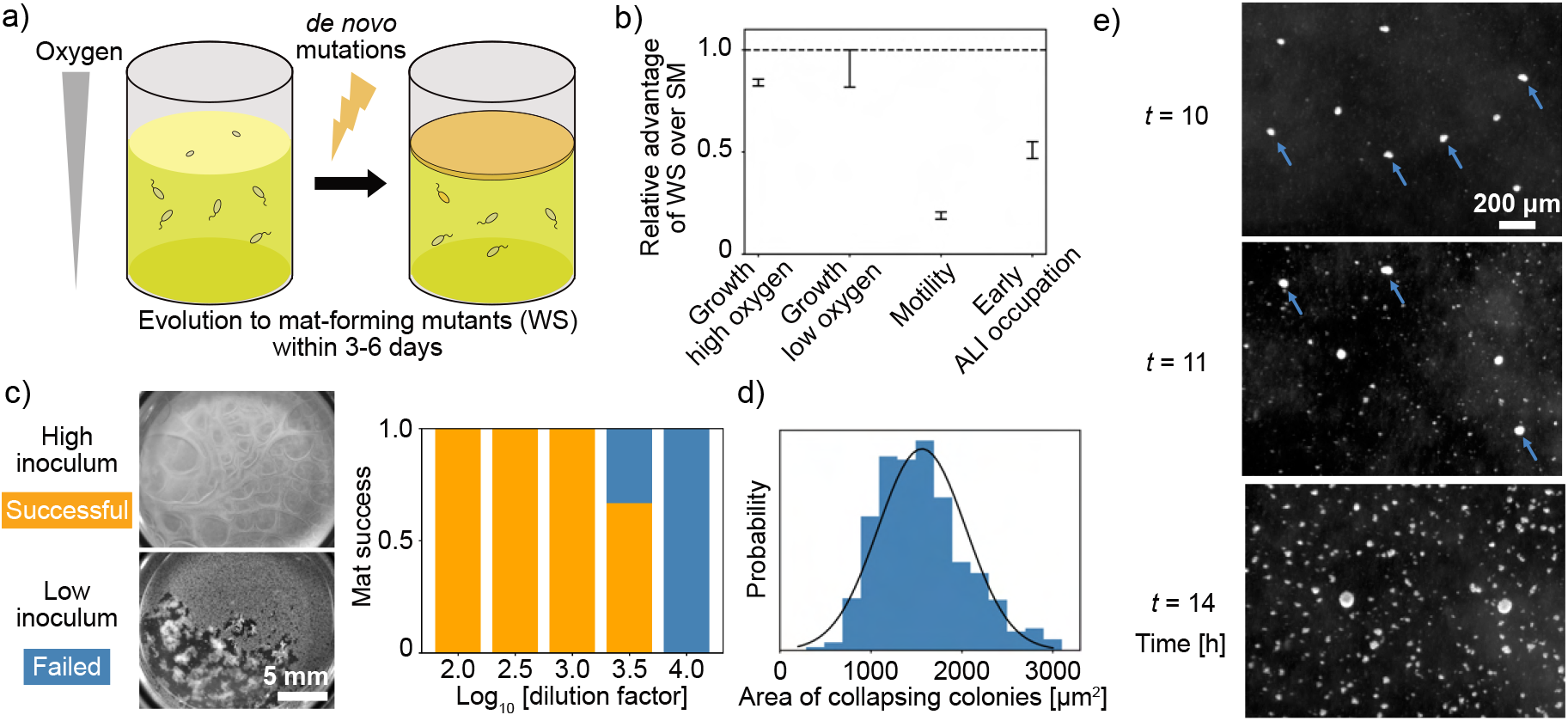
Adaptive mutants exhibit context-dependent fitness limitations. (a) *P. fluorescens* SBW25 reproducibly evolves mat-forming WS mutants in static culture within 3–6 days. (b) Summary of WS phenotypes compared to the ancestral SM type. The ratio of a quantified trait of WS over that of SM was plotted for each advantageous trait. Growth rate data are from Koza *et al*.^6^. Motility was measured using a semi-solid agar assay. Early ALI occupation was quantified as the ratio of surface cell density at *t* =2 h to the initial inoculum density. (c) WS mutants failed to form a mat in a monoculture when initiated from low inoculum densities. Mat formation success was scored at *t* =24 h across a dilution series. Organe bars show the ratio of successful populations, and blue bars how the ratio of failed populations. *N* =3, 6, 12, 15, 9 for each dilution factor (from 10^2^ to 10^4^). (d) Distribution of WS microcolony sizes immediately prior to collapse in monoculture. Black line indicates a Gaussian fit. (e) Dark-field images showing microcolony collapse events in WS monocultures. Blue arrows mark detached colonies falling from the ALI.

To quantify fitness costs during early ALI colonization, we measured the surface cell density at the ALI across a range of inoculum densities. A representative WS mutant (caused by deletion of *wspF*. See MPB08384 in SI table S3) showed lower surface densities than SM, indicating reduced colonization efficiency (Fig. 1b, “Early ALI occupation”). Similar trends were observed for other WS genotypes (*awsX* and *mwsR* mutants, SI table S3), although some showed slightly higher efficiency than SM (Fig. S1). None, however, displayed a clear advantage that could explain the rapid and reproducible evolution of WS.

A more striking result emerged when we further varied the inoculum density. WS mutants failed to form a mat when cultured alone from low inoculum densities (∼10^6^ cells/mL, Fig. 1c, Supplementary Movie 1). Although inoculum effects on biofilm formation have been reported previously^18,19^, complete mat failure at low density appears to have gone unnoticed. The failure occurred during the initial 24-hour colonization window and was not observed in subsequent rounds of mat formation, which proceeded successfully regardless of inoculum size. This finding raises an important question: how do spontaneous WS mutants in laboratory evolution colonize the ALI so reliably if they cannot establish from low density—their likely state upon first arising?

To explore this paradox, we used microscopy to track ALI microcolony formation at low inoculum. In WS monocultures, microcolonies repeatedly collapsed once they reached a diameter of approximately 40 µm, producing hole-like defects that ultimately prevented coherent mat formation (Fig. 1d,e, Supplementary Movie 3).

These collapses are consistent with a size-dependent physical instability of floating microcolonies. The mass densities of bacterial cells (∼1.1 g/cm^320,21^) and extracellular matrices (in the case of cellulose, it is ∼1.5 g/cm^3^ or ∼1.37 g/cm^3^ in an amorphous state^22,23^) exceed that of the surrounding medium. For small colonies below the capillary length (on the order of millimeters), surface tension provides the primary support. However, surface tension scales with length, while gravitational force scales with volume. As a result, microcolonies become unstable once they exceed a critical size, consistent with our observations.

To formalize this dynamic, we developed a spatial model simulating early ALI colonization (see SI for details). The model distinguishes between a well-mixed bulk phase and a structured ALI, represented as a two-dimensional lattice. Colonization proceeds via local growth and aggregation of surface-attached microcolonies. Successful mat formation requires that colonies connect to form a coherent mat before reaching the size threshold for collapse. The resulting dynamics were characterized by sharp shifts in success rate as a function of inoculum density (Fig. 2c,d), resembling a percolation transition^24^, in which the number of connected clusters increases drastically. Simulations also revealed that traits enhancing surface dispersal—such as motility—significantly increased colonization success (Fig. 2b,d), underscoring the disadvantage of WS mutants in this early phase.

**Figure 2.**
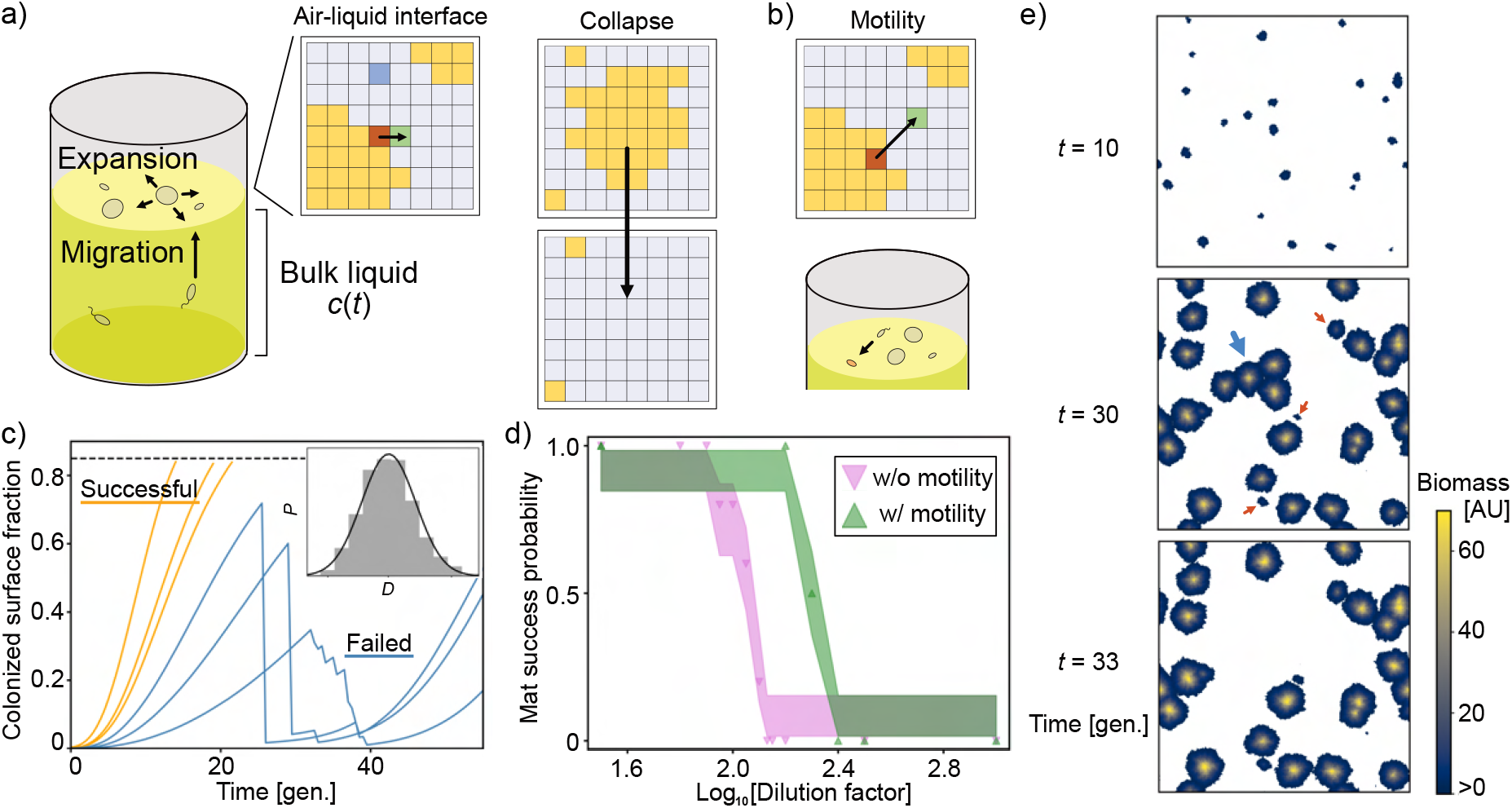
Spatial simulations reveal inoculum-dependent success in ALI colonization. (a, b) Schematics of the simulation model. The culture is divided into a well-mixed bulk liquid and a structured ALI represented as a 2D lattice. Microcolonies grow, merge, and collapse. Collapsing occurs based on size-dependent physical instability. See SI for model details. (c) Example simulation trajectories without motility. Mat formation was successful at higher inoculum densities (orange lines; dilution factors 10^1.5^ to 10^1.9^), but failed at lower inocula (blue lines; dilution factors 10^2.2^ to 10^3^). When 85 % of the ALI was covered (dotted line), simulations were terminated, and mat formation was regarded as successful. Inset shows size distribution of collapsing isolated colonies (*D*: diameter in an arbitrary unit. *P*: normalized frequency); solid line is a Gaussian fit. (d) Probability of successful mat formation by *t* =30 [generation] as a function of inoculum density, with (green) and without (pink) motility. Shaded regions show 68 % confidence intervals from a Beta distribution (*N* =10 replicates per condition). (e) Simulation snapshots at a low inoculum (dilution factor 10^3^). Early colonizers attach at *t* =10; later colonization occurs via migration from the bulk (red arrows at *t* =30). Large aggregates collapse due to instability (blue arrow at *t* =30).

### Compensation for WS fitness costs

The failure of WS mutants to form mats at low inoculum raises a fundamental question: what enables their repeated success in evolution experiments? A key difference between these monoculture assays and standard evolution protocols is the presence of the ancestral SM type. We hypothesized that SM cells might play a direct role in facilitating WS establishment.

To test this, we performed co-culture experiments. When SM cells were added to low-inoculum WS populations (conditions under which WS alone failed to form a mat), mat formation was restored in approximately 20 % of replicates (2 out of 9). Microscopy revealed that SM and WS cells co-colonized the ALI (Fig.3a). Based on insights from our spatial model (Fig.2), we infer that the presence of SM stabilizes early ALI colonization by helping to connect neighboring microcolonies, thereby preventing collapse. These results suggest a positive mechanical interaction between the two types, which has not been previously reported.

To investigate the nature of this interaction, we tracked microcolony growth in co-cultures using fluorescence microscopy. When mixed at equal frequency, WS and SM jointly colonized the ALI, and no clear differences in growth or expansion were observed (Fig. 3b). This indicates that the positive effect of SM on WS establishment is not due to a direct fitness gain for WS, but rather emerges from the physical properties of collective colonization.

**Figure 3.**
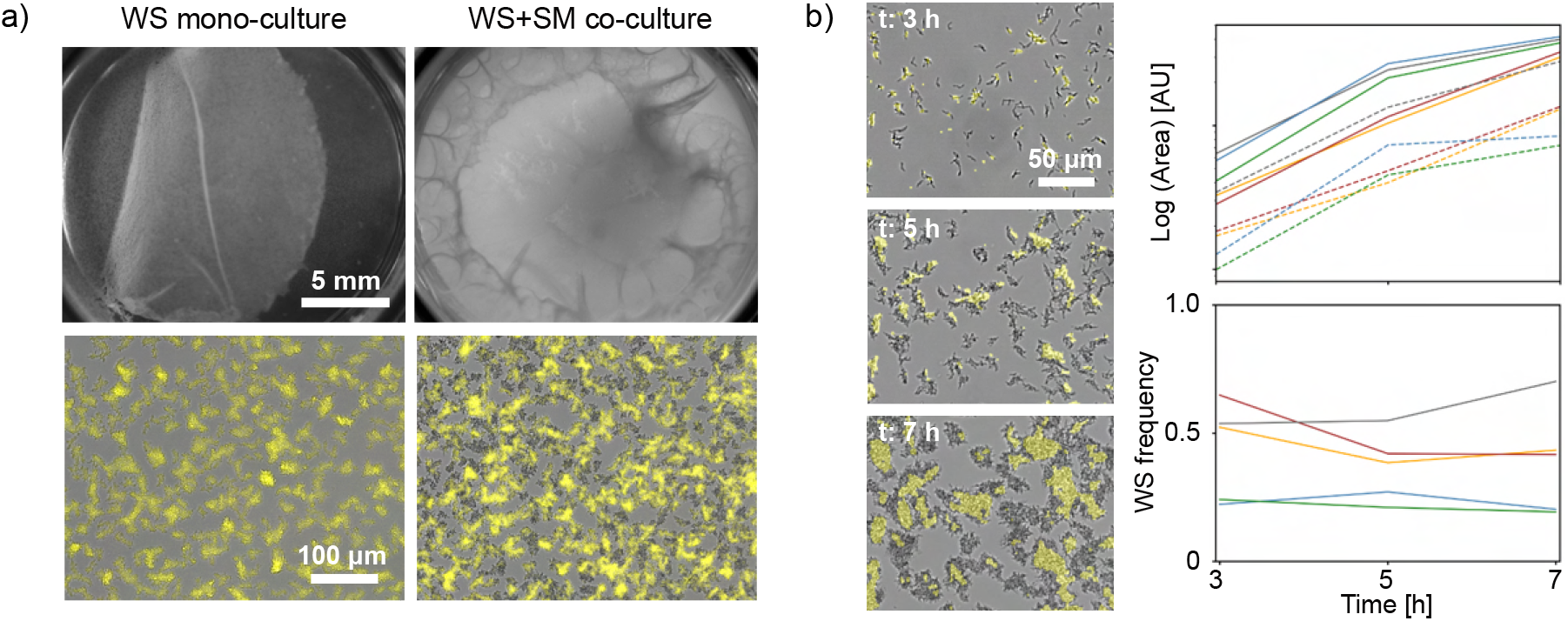
Ancestral SM cells enable WS colonization through mechanical facilitation. (a) Mat formation failed in WS monocultures at low inoculum, but was rescued in ∼20% of replicates (2 out of 9) when a similar amount of SM cells were co-inoculated. Bottom panels show ALI colonization at *t* =7.5 h, with WS visualized using a fluorescent marker. (b) Fluorescence time-lapse microscopy of WS–SM co-cultures revealed no clear differences in expansion rate or competitive fitness at the ALI. The top plot shows the area expansion of the total population (solid) and WS population (dotted). The change in WS frequency is shown in the bottom plot. Colors exhibit experimental replicas, while the fluorescent tag was flipped in blue and green replicas.

We next examined the role of ancestral SM cells in the evolutionary dynamics. To do this, we used a SM genotype carrying a c-di-GMP–responsive reporter (pCdrA–mScarlet)^25,26^. The reporter is non-fluorescent under basal conditions but becomes fluorescent when intracellular c-di-GMP levels rise, allowing tracking of the emergence and expansion of WS mutants, which over-produce c-di-GMP.

When the strain was cultured under static conditions, fluorescence was undetectable during early growth but became visible after the ALI was fully colonized by non-fluorescent SM cells. At *t* =24 h, we detected a small fluorescent colony at the ALI, and by *t* =48 h, multiple fluorescent microcolonies had appeared (Fig. 4a). These foci resembled locally growing WS clusters, consistent with their reduced motility and limited dispersal. To confirm that fluorescence reflected WS identity, we plated evolved ALI populations on agar plates. Multiple colony morphologies typical of WS were recovered, most showing fluorescence. In contrast, fluorescent SM colonies were rarely observed (Fig. 4b,c). The existence of dark WS and fluorescent SM colonies might be explained by the sensitivity of the reporter or exceptional correspondence between colony morphology and ALI colonization^26,27^.

**Figure 4.**
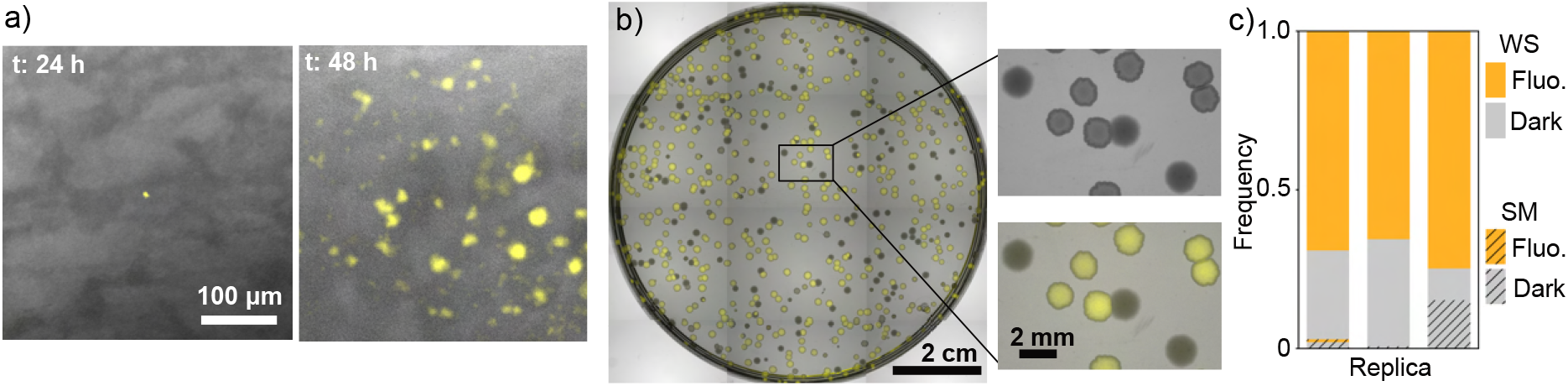
Spatial dynamics of WS emergence revealed by a c-di-GMP reporter. (a) Ancestral SM cells carrying a pCdrA–mScarlet reporter were evolved in static culture. Fluorescence was undetectable early on but appeared around *t* =24 h, after the ALI was fully colonized by non-fluorescent cells (*t*∼ 10 h). By *t* =48 h, multiple fluorescent microcolonies were visible across the ALI. (b) After six days of evolution, cells from the ALI were plated on agar plates. WS and SM types were distinguished by colony morphology. Fluorescence was primarily observed in WS colonies. (c) Frequency of colony morphotypes and fluorescence expression across replicate lines. More than 1000 colonies were scored per treatment.

Together, the c-di-GMP reporter experiments reveal the spatio-temporal sequence of WS evolution: (i) SM transiently colonizes the ALI, (ii) WS mutants emerge locally within the SM population at the ALI, and (iii) WS expands via local growth. Pre-colonization by SM appears to act as a physical scaffold that enables newly emerging WS cells to remain surface-associated and avoid microcolony collapse. Moreover, the presence of SM at the ALI increases the likelihood of WS mutations arising directly at the interface. This spatial positioning offsets the motility disadvantage of WS mutants, as they no longer need to migrate from the bulk to access the niche. These results highlight the central role of ancestral populations in facilitating adaptive evolution of WS phenotypes.

### How slow-growing WS outcompetes SM

While our results show that SM cells can compensate for the fitness costs of WS mutants during colonization, this alone does not explain how WS types ultimately increase in frequency and dominate the ALI population. Given that WS mutants grow more slowly than SM, what allows their eventual dominance?

We hypothesized that the answer lies in the general transition between sessility and dispersal in biofilm-forming bacteria^8^. Many bacterial species regulate motility and extracellular matrix production in response to environmental conditions, often using c-di-GMP as a key signaling molecule. During biofilm development, cells frequently shift to a motile state to disperse once the biofilm matures^28^. In *P. fluorescens*, WS mutations elevate c-di-GMP levels^16^, locking cells into a sessile state and reducing their capacity to disperse.

If SM cells are more prone to dispersal, then WS may gain an indirect advantage simply by remaining localised at the ALI (Fig. 5a). To test this, we tracked motility transitions in a SM-majority population. Initially, SM cells formed a weak mat that covered the ALI by *t* =10 h. Shortly after, many cells became motile, leading to phase separation between active fluid regions and sessile microcolonies (see Supplementary Movie 4). Using particle image velocimetry (PIV), we observed dynamic fluid motion within these motile regions over a 60-minute interval (Fig. 5b). In contrast, WS clusters remained largely sessile and stable.

**Figure 5.**
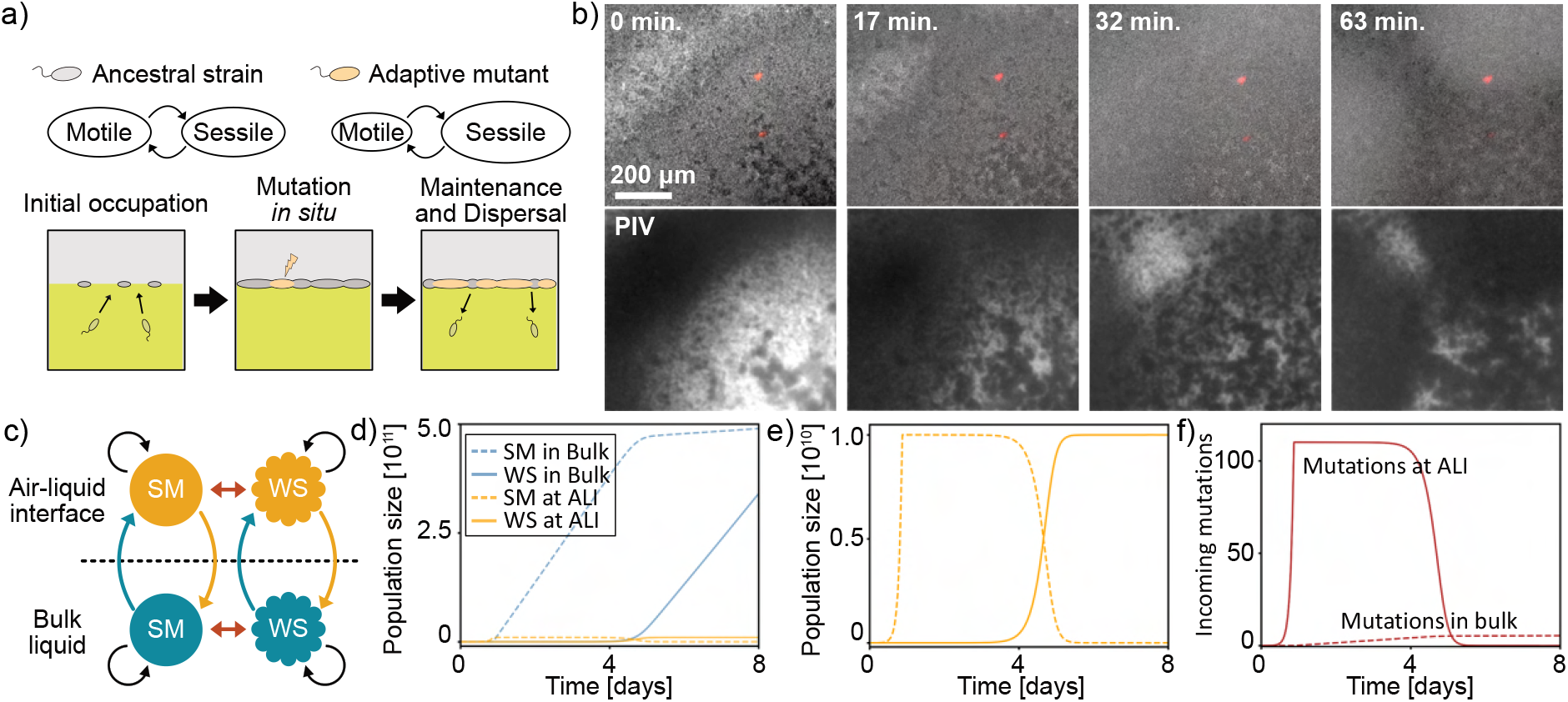
Dispersal-driven dynamics enable WS takeover despite growth disadvantages. (a) A conceptual schematic of the dispersal-driven takeover. SM cells exhibit greater motility and are more likely to disperse from the ALI, while WS cells remain sessile and persist at the interface. (b) Transition to motility in an SM-majority population. By *t* =11 h, active fluid regions emerged as SM cells became motile. Bottom panels show time-averaged velocity magnitudes (|*v*|) over 5-second intervals. WS clusters (in red) remained stable and sessile. (c) A schematic of a four-state model with two genotypes (SM and WS) and two spatial environments (ALI and bulk liquid). The four states develop via growth (black arrow), migration from bulk to ALI (blue arrow), dispersal from ALI to bulk (orange arrow), and mutational transition (red arrow). WS has lower growth and migration rates than SM but benefits from reduced dispersal from the ALI. (d) Simulated population dynamics of each state over time. Despite lower intrinsic fitness, WS takes over due to higher retention at the ALI. (e) Magnified population dynamics at the ALI. (f) The number of incoming mutations (from SM to WS) at the ALI increases steeply over time, reflecting the local emergence and coexistence of multiple adaptive WS lineages.

To formalize this dynamic, we developed a four-state model incorporating two cell types (SM and WS) and two spatial locations (ALI and bulk liquid) (Fig. 5c). The population size of each state changed over time via growth, spatial replacement, and mutation of cells. WS was assigned lower growth and migration rates than SM, reflecting its fitness disadvantages. Its sole advantage was a reduced tendency to disperse from the ALI. Assuming differential oxygen availability between the ALI and the bulk, and a carrying capacity constraint at the interface (see SI for details), the model reproduced WS takeover, despite disadvantages to individual cells (Fig. 5d,e).

The model simulations are consistent with our c-di-GMP reporter experiment (Fig. 4), showing the in-situ emergence and expansion of the WS type at the ALI upon SM pre-colonization (Fig. 5e,f). Notably, the model also predicts a steep increase in the number of newly arising mutations at the ALI, suggesting the emergence and coexistence of multiple adaptive WS lineages (Fig. 5f).

### Diversity of WS lineages in a single evolved population

The evolutionary dynamics of WS mutants in static cultures depart markedly from the classical selective sweep model, in which a single high-fitness lineage fixes in the population. WS mutants exhibit multiple fitness disadvantages and depend on the presence of ancestral SM cells to establish at the ALI. Their spread is driven less by intrinsic growth rate than by reduced dispersal, suggesting that colonization success may not require competitive exclusion.

These features raise the possibility that multiple WS lineages could emerge and coexist within a single evolving population. To test this, we performed evolution experiments using a 1:1 mixture of two SM genotypes, each carrying a different fluorescent tag. After six days of static culture, the resulting mat populations were plated on agar plates, and individual colonies were scored for both color and morphology (Fig. 6a).

**Figure 6.**
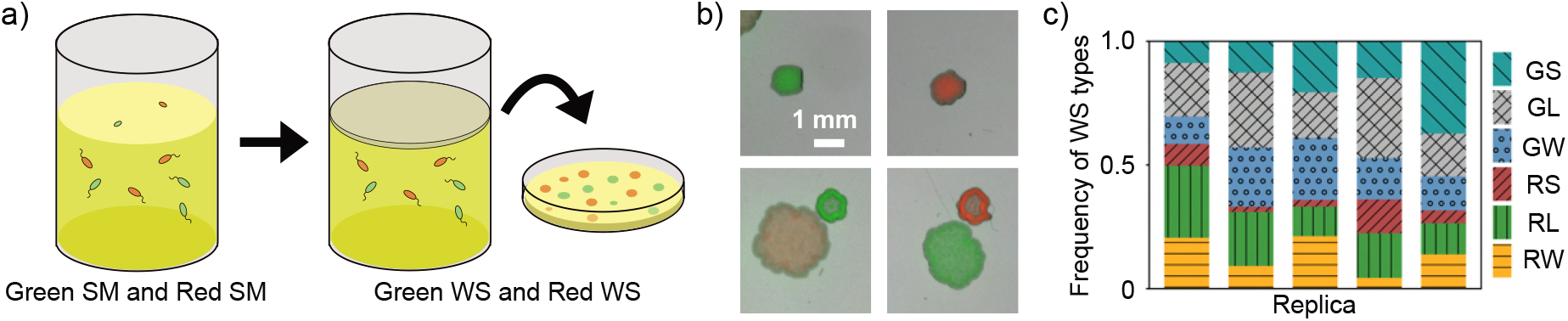
Coexistence of diverse WS lineages within evolved populations. (a) Schematic of the experimental setup. Two ancestral SM genotypes, each tagged with a distinct fluorescent marker (green and red), were mixed 1:1 and evolved under static conditions for six days. The resulting mat populations were plated on agar plates to assess colony color and morphology. (b) Representative image showing the six observed WS colony types, defined by combinations of color and morphology. (c) Distribution of WS morphotypes across five independent evolution lines. More than 140 colonies were scored per replicate. All six morphotypes were consistently recovered, indicating the co-emergence and coexistence of multiple WS lineages within single evolved populations. Letters show colors (G: green, R: red) and morphs (S: small WS, L: large WS, W: wheel-like WS).

WS colony morphologies were classified into three types—small, large, and wheel-like—following established criteria^29^. Across five replicate evolution lines, all six combinations of color and morph (2 colors × 3 morphs) were consistently observed (Fig. 6b,c), confirming the presence of multiple distinct WS lineages within each population. Since different genotypes can produce similar morphologies, the true genotypic diversity of WS mutants is likely even higher.

While a full analysis of WS diversity and its scaling with population size is beyond the scope of this study, such patterns could be explored using approaches such as deep sequencing or genetic barcoding, which have recently been developed for *P. fluorescens* SBW25^30^. Our results nonetheless demonstrate that WS diversification is a consistent outcome of evolution in this system, shaped by spatial structure, ecological context, and dispersal dynamics.

## Discussion

The consistent emergence of wrinkly spreader (WS) mutants in static cultures of *P. fluorescens* SBW25 has long served as a model for studying adaptive radiation and biofilm formation^1^. Despite their reproducible appearance, WS mutants possess inherent disadvantages, such as reduced growth rates^6,11^and impaired motility^12,16^, raising questions about their evolutionary success. Previous work has attributed success to strain-specific properties of the mutants, such as over-production of cellulose^31^ and surfactant secretion^12^. Here, our study highlights the importance of ecological context^32–34^, revealing that prior colonization of the air-liquid interface (ALI) by ancestral smooth (SM) cells creates a scaffold that facilitates the establishment and eventual dominance of WS mutants.

This “dispersal-driven” dynamic underscores a broader ecological principle: transition between motility and sessility in biofilm-forming bacteria^8^. In typical biofilm life cycles, cells switch between motile and sessile states in response to environmental conditions^28^. WS mutations, characterized by elevated c-di-GMP levels, lock cells into a sessile state, which, while disadvantageous in fluctuating environments, becomes beneficial in the stable niche of the ALI. This explains the consistent emergence of WS mutants in laboratory settings.

Our findings also highlight a novel eco-evolutionary interaction mediated by a mechanical effect. The pre-colonization of the ALI by SM cells provides a physical scaffold that enables the establishment of WS mutants. This interaction is reminiscent of the evolution of auxotrophs, where ancestral cells produce metabolites essential for mutant survival^35,36^. However, in our system, the interaction is purely mechanical, arising from the trade-off between extracellular matrix production and motility. Such mechanical interactions may be widespread in mat-forming bacteria across various species and environments.

Two features of our model system warrant consideration. First, the oxygen gradient between the bulk liquid and the ALI is fundamental. SM occupation of the ALI prevents oxygen supply to the bulk, suppressing the growth and viability of mutants arising there. Consequently, in-situ mutations at the ALI are more likely to succeed. Moreover, WS mutants, by virtue of our discoveries here, must in fact arise at the ALI and not migrate to this niche from the bulk phase. Second, the relatively large population size at the ALI (∼10^10^ CFUs for ∼400 mm^2^ ALI area) increases the probability of beneficial mutations occurring *in situ*. In smaller niches, such as colonic crypts in mammalian guts, colonization by mutants may rely more on rare migration events from external populations.

The question of whether migration or mutation occurs first in niche colonization is pertinent. Given the trade-off between motility and adhesion, mutations that enhance niche colonization often impair dispersal. Our study demonstrates that pre-colonization by ancestral cells can facilitate the establishment of such mutants. However, in scenarios where ancestors cannot pre-colonize the niche, two evolutionary pathways are conceivable: (i) a one-step mutation enables niche colonization followed by rare migration, or (ii) an initial mutation enhances motility, allowing niche occupation, followed by a second mutation that promotes sessility. The latter pathway may be more likely when migration is particularly challenging. Indeed, laboratory experiments have shown the evolution of niche colonizers from ancestors lacking key components for ALI colonization, such as strains deficient in c-di-GMP pathways (Δ*wsp*Δ*aws*Δ*mws*)^37,38^ or cellulose production (Δ*wss*)^39^. These evolved types often exhibit reduced motility, with mutations affecting chemotaxis or flagellar function^16,38^. Such observations support the plausibility of a two-step evolutionary process in niche colonization.

In summary, our findings reveal that the reproducible emergence of WS mutants is not solely a consequence of their intrinsic properties but is significantly influenced by the ecological context established by ancestral SM cells. This underscores the importance of considering eco-evolutionary dynamics and physical interactions in understanding adaptive evolution.

## Methods

### Strains and culture conditions

*Pseudomonas fluorescens* SBW25 strains (see SI table S3 for the strain list) were cultured in King’s B medium (KB)^40^ at 28 ^*°*^C. To avoid the strong autofluorescence of KB in the green wavelength, mScarlet was used as the main fluorescent marker for bacteria. A *wspF* mutant^41^ was used as a typical WS type. Prior to experiments, cells were streaked on an agar plate from a frozen stock and incubated for 2 days. To obtain a saturated cell suspension, cells from a single colony on a plate were taken and cultured in a liquid medium with continuous shaking for 24 hours. When diluting or transferring cells, cell suspensions were agitated well with a vortex mixer for more than one minute to resolve clumps.

### Measurements of motility and early ALI occupation

Swimming motility was quantified on freshly prepared semi-solid agar plates (0.25% agar and 1% KB, SSA) following the previously established method^26^. 200 µL of 24-hour-old shaken liquid cultures was transferred to wells in a 96-well plate after brief vortexing. The cell suspensions were set down for ∼1 hour to let debris settle, and cells were carefully sampled with a toothpick from the top surface of the liquid. The toothpick was vertically stabbed in the center of an SSA plate. SSA plates were incubated at 28 ^*°*^C for 24 hours. The swimming halo diameter was measured with a digital vernier (Caliper).

To quantify the early ALI occupation, cells at the ALI were observed by an upright microscope (Axio Imager Z2, Zeiss) with a 10x objective closely placed above the ALI. Starting cultures were inoculated in a 12-well microtiter plate (TPP) and incubated in a microscope incubator (PeCon). To measure the surface density, the number of cells at *t* =2 h was directly counted around the center of the ALI, where the effect of the air-liquid meniscus on the z focus is minimized. The measurement was iterated across various volume inoculum densities. The volume densities were quantified by plating diluted suspensions on agar plates and counting the CFUs. The ratio of the surface density to the inoculum density was defined as a measure of the efficiency of early ALI occupation.

### Observation of mat formation and surface colonization

A saturated liquid pre-culture was diluted to the required inoculum densities in a 12-well microtiter plate (TPP) to start static cultures. The macroscopic dynamics of mat formation were tracked by dark-field imaging with a stereo microscope (Axio Zoom V16) every 10 minutes for 24 hours. During static cultures, a stage-top incubator with a heated lid (ibidi) was used to reduce evaporation. The timing of the mat collapse was manually detected from timelapse images. To analyze the critical size for a single colony to collapse, a magnified dark-field image (140x) was taken at the center of an ALI, and the colony size at the time immediately prior to collapse was measured. The colony size was quantified as the area of connected pixels after applying the Otsu binarization and closing morphological operation. To observe the microscopic configuration of microclonies at the ALI, an upright fluorescence microscope (Axio Imager Z2, Zeiss) was employed. To take a timelapse, XY drift due to strong surface fluctuations of the ALI was compensated by manual XY adjustments.

### Evolution experiments

To track the spatio-temporal course of evolutionary dynamics, an SM strain carrying a c-di-GMP-responsive reporter (pCrA-mScarlet) was cultured from a high inoculum density (∼10^7^ cells/mL) for 6 days. 50 µg/mL Gentamicin was added to the culture media to maintain the plasmid and prevent contamination. The spatial expression of fluorescence was tracked with an upright microscope (Axio Imager Z2). To examine the fluorescence and the morphotype of evolved cells, the mat structure was carefully sampled from the ALI with a pipette. The suspension was centrifuged at a low speed (150 rcf) for 2 minutes (MiniSpin, Eppendorf). The supernatant was gently removed, and the sediment was resuspended in 1 mL PBS. The sample was sonicated for 15 seconds (Sonopuls mini20, Bandelin) to resolve the mat structure and plated at 10^6^ and 10^7^ dilutions. The colonies on plates were manually scored with a stereo microscope (Axio Zoom V16).

To evaluate the diversity of WS lineages in an evolved population, a 1:1 mixture of SM strains carrying different fluorescent tags (sGFP2 and mScarlet) was statically cultured from a high inoculum (∼10^7^ cells/mL) at 28 ^*°*^C. At six days, the evolved mat population was sampled and plated as described above. The fluorescence and the morphology of colonies were manually scored by a stereo microscope (Axio Zoom V16). The colony morph of WS was classified into three subclasses following the established criteria ^29^.

### Observation of motility transition at the ALI

To track the transition to motility at the ALI, a small number (∼10^3^ cells/mL) of WS cells carrying a fluorescent tag (mScarlet) were mixed into an SM-majority (∼10^6^ cells/mL) population. The suspension was cultured under a microscope incubator, and the spatio-temporal dynamics were tracked with an upright microscope (Axio Imager Z2). At each time point, after manually correcting XY drift, a snapshot of fluorescence expression was taken, and the active movements were recorded for 5 seconds with bright-field imaging. To access the local activity, the XY drift was computationally corrected based on the feature pattern of an arbitrarily-chosen sessile region, and particle image velocimetry (PIV) was conducted with PIVlab^42^ in MATLAB (MathWorks). The XY drift was further corrected by subtracting the spatially-averaged velocity of a sessile region. The magnitude of the velocity at each position was averaged over 5 seconds to quantify the local activity.

### Model simulations

The early-phase colonization at the ALI was simulated with a spatial model that distinguishes between a well-mixed bulk phase and a structured ALI phase, represented as a two-dimensional lattice (see SI for details). Cells in the bulk liquid grow logistically and migrate to the ALI to colonize a random vacant position at a rate proportional to the bulk population size. At the ALI, colonies grow horizontally following the Eden model rule^43^ as well as accumulate biomass locally. The stability of an ALI colony is evaluated by the net biomass and the colony periphery length, and unstable colonies are removed from the ALI. The stochastic simulations are conducted by the next reaction method^44,45^and terminated once 85 % (chosen arbitrarily) of the ALI is covered by cells.

To formalize “dispersal-driven” evolutionary dynamics, we developed a four-state model (see SI for details) with two cell types (SM and WS) and two spatial locations (bulk liquid and ALI). The time development of the four states is driven by growth, spatial relocation, and genetic mutations. We assumed a carrying capacity of an ALI reflecting a spatial constraint (the finite ALI area). With this constraint, the dispersal rate from the ALI and migration rate to the ALI are assumed to be dependent on the ALI population size. To reflect the strain fitness, WS is assigned lower growth and migration rates than SM. The sole positive trait of WS is reduced dispersal rates from the ALI. The oxygen depletion is implemented as a reduction in the growth of both cell types in the bulk liquid. Simulations are started from SM-only populations.

## Data availability

The Supplementary Movies, spreadsheet data, and scripts are available at Zenodo (https://doi.org/10.5281/zenodo.15293177). The microscopy data are available at BioImage Archive (https://doi.org/10.6019/S-BIAD2117).

## Acknowledgement

We thank Andrew D. Farr, David Rogers, and Loukas Theodosiou for helpful discussion and comments, and Carsten Fortmann-Grote for advice on employing computational clusters and organizing the data repository. We acknowledge Ellen McConnell, Joanna Summers, Lena Meister, and Michael Barnett for constructing the strains in the Microbial Population Biology strain repository at the Department of Microbial Population Biology, Max Planck Institute for Evolutionary Biology, Germany. This work was enabled by generous core support from the Max Planck Society.

## Supplementary Information

### Supplementary Methods

#### Confocal imaging and 3D reconstruction of microcolonies

To observe the 3D structure of an ALI microcolony, a *wspF* WS mutant with a fluorescent tag (sGFP2) was cultured in a 35 mm plastic dish (Sarstedt) and observed with an upright confocal microscope (LSM880, Zeiss). To reduce the autofluorescence, lysogeny broth (LB) was used instead of KB. Moreover, *pvdS*, a gene necessary for expression of the fluorescent siderophore pyoverdin, was deleted in order to eliminate the pyoverdin-generated fluorescence signal. Static culture was started from a low inoculum (∼ 10^4^ cells/mL) and observed after 13 hours of static incubation. Colonies at the ALI were captured with a 10X objective, closely located above the ALI. To capture rafting colonies, the speed of image acquisition was increased to minimize drift, albeit at the expense of image quality. The z-stacks of the confocal images were reconstructed by the 3D viewer of ImageJ.

### Co-culture experiments to capture the spatial configuration of SM and WS

To capture the interactions between SM and WS at a micron scale, *pvdS* was deleted from bacteria, and LB, instead of KB, was used as the culture medium to reduce the autofluorescence. SM and WS (*wspF* mutant) carrying fluorescent markers (sGFP2 and mScarlet) were co-cultured for 16 hours under static conditions from a low inoculum (∼ 10^3^ cells/mL). The spatial configuration of the two types was captured with an upright microscope (Axio Imager Z2, Zeiss).

## Supplementary Texts

### A Stability condition for floating microcolonies at an ALI

Things smaller than the capillary length of liquid can float at the air-liquid interface even if they are heavier than the liquid. Two lifting forces, buoyancy and surface tension, counter a gravitational fall. The gravitational force can be expressed by the mass density of a colony *ρ*_*c*_, the volume of the colony *V*, and the gravitational acceleration *g*,

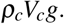

The buoyancy term is proportional to the mass of the displaced liquid by the colony. Letting the displaced liquid volume be *V*_*below*_, the buoyant force is

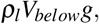

where *ρ*_*l*_ is the mass density of the liquid. Here, we assume *ρ*_*l*_ *< ρ*_*c*_ and *V*_*below*_ *< V*_*c*_. The lifting force by surface tension works at the air-liquid-colony interface, and can be calculated by a line integral across the periphery:

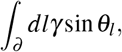

where ∫_∂_ *dl* is an integral over the periphery, *γ* is the surface tension of the liquid, and *θ*_*l*_ is the tilting angle of the deformed liquid surface near the colony. In general, *θ*_*l*_ is very small and sensitive to the local properties, such as the shape or the hydrophobicity of the colony. Here, for simplicity, we assume a constant *θ* across the periphery and get

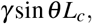

where *L*_*c*_ is the colony periphery length. The force balance equation of a floating colony is

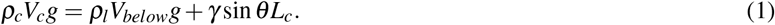

For simplicity, we assumed that *V*_*below*_ was proportional to *V*_*c*_:

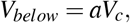

where *a*(*<* 1) is a constant. The equation 1 can be simplified as

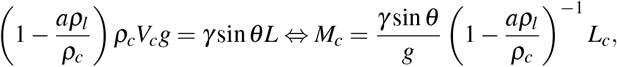

where *M*_*c*_ =*ρ*_*c*_*V*_*c*_ is the total biomass of the colony. By defining

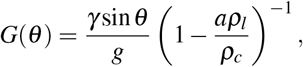

we get the simplified force balance equation

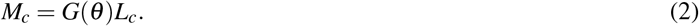

As a colony gets bigger, the left-hand side grows as ∼*O*(*L*^3^), while the right-hand side grows as ∼*O*(*L*^1^). Therefore, when the colony gets too big, no *θ* can satisfy the equation 2, and the colony falls down from the air-liquid interface.

In computational simulations, we fixed *θ* regardless of the colony shape and size, and defined a constant *G* to evaluate the stability of a colony.

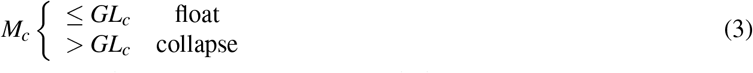

We empirically chose *G* to let an isolated colony collapse at the colony size consistent with the experimental result (Fig. 1).

### B Simulations of early-stage spatial dynamics of ALI colonization

Static microbial culture is modeled as a combination of two populations, the well-mixed population in the bulk liquid phase (supposedly mixed by hydrodynamic convection, often driven by biological activities^13,46^) and the spatial ALI population on a two-dimensional lattice. The population density in the liquid phase *c*(*t*)?is modeled as a logistic growth with the growth rate *r*_*l*_ and the carrying capacity *c*_*max*_.

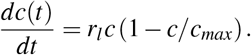

Note that *r*_*l*_ should be smaller than that of a shaken culture because of oxygen depletion. The starting cell density *c*_0_ is varied in the simulation to investigate inoculum effects. From the liquid phase, cells migrate to the air-liquid interface and colonize a random position at migration flux *µ*, which is proportional to *c*(*t*):

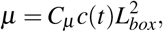

where *C*_*µ*_ is a coefficient, and 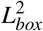 is the entire area of the air-liquid interface.

The ALI population is modeled on a two-dimensional discrete space with a periodic boundary condition. Once a cell occupies a site, the cellular colony grows in the horizontal (spatial expansion) and vertical (local biomass accumulation) directions. The horizontal expansion follows the Eden-model-type surface growth^43^: only the cells facing to an empty space can expand and colonize a neighbor site.

To model the local biomass accumulation, we referred to the three-dimensional structure of a colony on a plate. It has been known that the height of a colony saturates due to the limited nutrient diffusion^47^, and the saturation can be modeled by a logistic growth^48^. Thus, the biomass at a site (*x, y*), *M*(*x, y, t*), follows

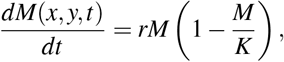

where *r* is the vertical growth rate and *K* is the maximum local biomass. Note that intra-colony displacements of biomass, such as the diffusion term *D*∇^2^*M*, are not incorporated into the model for simplicity, and thus the biomass grows only locally. During the simulation, the stability of colonies (eq. 3) is tested at each time point, and unstable colonies are removed.

We also implemented the motility of cells as a stochastic replacement from a colony periphery to another random site. The replacement probability is defined by the rate *r*_*s*_. We set *r*_*s*_ =0 for non-motile simulations.

To conduct simulations, the modified next reaction method^44,45^is used. Note that the well-known Gillespie algorithm is not simply applicable to our model because we have the time-dependent variable *c*(*t*), which defines the propensity of migration from the bulk liquid to the air-liquid interface. Fortunately, the logistic function *c*(*t*) enables us to analytically treat the propensity in the method. In short, the logistically growing population

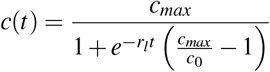

defines the propensity of migration, and the expected time to the next event Δ*t* can be calculated by

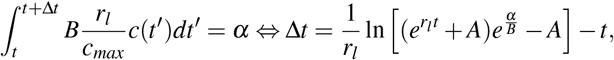

where *A* and *B* are constant, and *α* is related to the “internal time” of the next reaction method. Refer to^45^ for further details of the method. The step-by-step description of the model simulations is as follows:

#### Detailed algorithm

The model has three stochastic processes: Eden-model-type colony expansion (*g*), migration from the bulk liquid to the ALI (*m*), and cell replacement due to swimming motility (*s*). We label the processes by index *k* that can take *g, m, s* . The propensity of each process *a*_*k*_(*t*)?can be expressed as

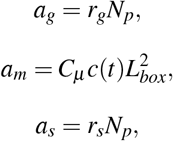

where *N*_*p*_ is the total number of periphery sites. We set the timescale of the simulation to let *r*_*g*_ =1. We define the internal time of each stochastic process *T*_*k*_ and the expected timing of the process in the internal time *P*_*k*_. With this setting, the expected timing of the next process in the real time Δ*t*_*k*_ is a solution of

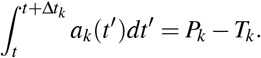

Note that for *k ∈* {*g, s* }, *a*_*k*_ is constant given no process happens during Δ*t*_*k*_, and the equation above can be easily solved. For *k* =*m*, fortunately, a logistic function allows analytic treatments, and no numerical analysis is required. The step-by-step algorithm of the simulation is:

1. Initialize the system. Set the initial population size of the bulk liquid and the ALI. Set *t* =0 and *P*_*k*_ =*T*_*k*_ =0.
2. Generate independent random numbers *s*_*k*_ from a uniform distribution over (0, 1). Set 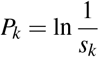 .
3. Calculate Δ*t*_*k*_ as the solution of 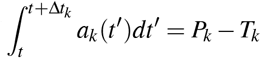
4. Set the actual time increment Δ =min_*k*_ {Δ*t*_*k*_} . If Δ ≤ *T*_*mass*_, let *k*^*′*^ be the process giving the minimum Δ*t*_*k*_. If Δ *> T*_*mass*_, set Δ =*T*_*mass*_, and no process happens except for the biomass growth (*k*^*′*^ =*φ*).
5. Set 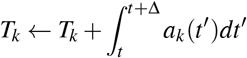, and then set *t*← *t* Δ.
6. Update the biomass of each site. And then update the site occupations based on the process *k*^*′*^ (if *k*^*′*^*≠φ*).
  - For *k*^*′*^ =*g*: Choose a random periphery site, and occupy one empty neighbor site.
  - For *k*^*′*^ =*m*: Occupy a random site. If the site is already occupied, the migration fails. For *k*^*′*^ =*s*: Choose a random periphery site (*x, y*). Draw a random number *d* from a Gaussian distribution with zero mean and variance *σ* ^2^. Scan the 3*σ* × 3*σ* neighbors of (*x, y*)?to find an empty site at a distance closest to |*d*| . If there are multiple candidates, choose one randomly. Colonize the new site with biomass *m*_0_. A swimming event fails if *d <* 0.5. We referred to the scheme in the previous literature^49^ but modified it to increase the computational efficiency.
7. Check the stability of colonies. Remove unstable colonies.
8. For *k*^*′*^, generate a new random number *s*_*k*_*′* from a uniform distribution over (0, 1), and set 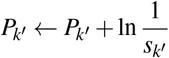.
9. Iterate the steps 3-8 until the terminating condition is reached, Note that this model was designed for conceptualizing the early-stage dynamics of mat formation to investigate the inoculum effect. To form a robust mat structure, strong adhesion to the side walls or mechanical cell-cell inter-connections by amyloid fibers should be required. We did not incorporate these additional complexities into our model.

### C Simulations of dispersal-driven evolution

To simulate evolutionary dynamics in an unshaken culture, we propose a simple four-state model. The model has 4 variables *S*_*A*_, *S*_*B*_,*W*_*A*_,*W*_*B*_, representing the population sizes of two cell types (SM and WS) and two spatial environments (bulk liquid and ALI). The simplest equations for the time development of the 4 states may be

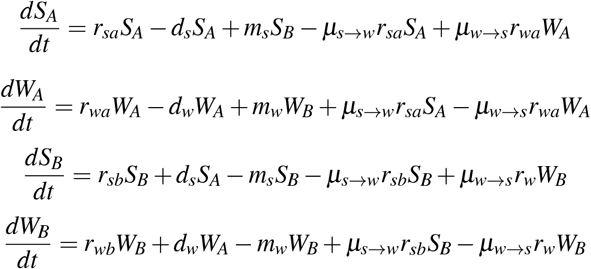

where *r, d, m*, and *µ* show the growth rate of each state, the dispersal of a strain from the ALI to the bulk, the migration of a strain from the bulk to the ALI, and the rate of mutation, respectively.

Here, we impose two constraints on the model to reflect the experimental setting:

1. The ALI population has a carrying capacity defined by the available space at the ALI.
2. Growth is not restricted by nutrients except for oxygen.

The second constraint is based on the fact that the growth in the bulk liquid is strongly reduced by oxygen depletion^6^. Therefore, the consumption of non-oxygen nutrients is very slow in the bulk liquid, and nutrients are continuously supplied to the ALI population by diffusion from the bulk. Considering the sufficient volume (5 ∼6 mL) of a liquid medium in evolution experiments, we assume that non-oxygen nutrients are not limiting the growth throughout experiments. As the oxygen gradient can be rapidly generated (within 4 hours^6^), we implement the effect of oxygen depletion into the constant parameters for the growth rates (*r*_*sa*_, *r*_*wa*_ ≫*r*_*sb*_, *r*_*wb*_).

With these constraints, the population size of the entire system is developed via the growth of the ALI population, whose size is upper-bounded by the spatial carrying capacity. Therefore, the entire system is supposed to grow almost linearly once the ALI population is saturated: cells are continuously produced at the ALI and transferred to the bulk liquid.

The spatial carrying capacity also means that the dispersal rate from the ALI to the bulk increases as the ALI population gets dense. We assume the crowding effect on the migration rate from the bulk to the ALI: fewer cells can move from the bulk to the ALI when the ALI population is spatially occupied. To this end, we consider the crowding-induced dispersal once the ALI population (*S*_*A*_ +?*W*_*A*_) reaches the spatial carrying capacity (*K*_*A*_).

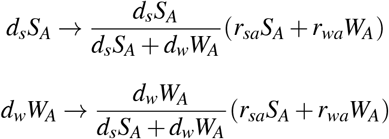

In addition, we limit the migration from the bulk liquid to the ALI by

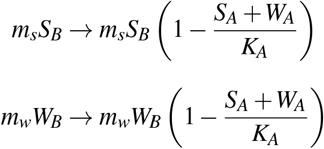

Note that the growth terms are unchanged. Here, the strain competition at the ALI is mainly driven by the ratio of the dispersal *d*_*s*_*S*_*A*_*/d*_*w*_*W*_*A*_.

The parameters are set to represent the experimental situations. Namely, *r*_*sa*_, *r*_*wa*_ ≫ *r*_*sb*_, *r*_*wb*_ (oxygen depletion), *r*_*sa*_ *> r*_*wa*_, *r*_*sb*_ *> r*_*wb*_ (slower growth of WS than SM), *m*_*s*_ *> m*_*w*_, and *d*_*s*_ *> d*_*w*_ (diminished motility of WS). In terms of the strain competition at the ALI, only the *d*_*s*_ *> d*_*w*_ condition (less dispersal) is regarded as the advantage for WS. In the model, we assume *d*_*s*_ and *d*_*w*_ to be constant for simplicity. If we increase *d*_*s*_*/d*_*w*_ upon the saturation of the ALI population to represent the difference in the motile-sessile switch, evolution will be accelerated.**z**

## Supplementary tables

**Table S1.** List of parameters in the spatial model simulations.

| Parameter | Explanation | Value |
| --- | --- | --- |
| $c_{max}$ | The maximum cell density in the bulk liquid. $10^9$ [cells/mL] | 5.0 |
| $r_l$ | The growth rate in the bulk liquid. | 0.15 |
| $r$ | The vertical growth rate of a colony at the air-liquid interface. | 0.2 |
| $m_0$ | The initial biomass for a newly colonized site. | 1.0 |
| $K$ | The maximum local biomass of a colony at the air-liquid interface. | 80 |
| $G$ | The coefficient for the stability of a colony. | 300 |
| $R_{end}$ | The ratio of colonized sites of the air-liquid interface to end a simulation. | 0.85 |
| $C_\mu$ | The coefficient to relate $c(t)$ and the migration rate $\mu$. | $10^{-5}$ |
| $L_{box}$ | The system size of the air-liquid interface. $L_{box} \times L_{box}$ was the entire 2D space. | 500 |
| $r_s$ | The rate of swimming events. | 0 or 0.05 |
| $\sigma$ | The spreading of the swimming distance. | 5.0 |
| $T_{mass}$ | The minimum time step for the mass growth and the stability evaluation. | 0.1 |
| $\varepsilon$ | The relative matrix production of the motile non-biofilm former. | 0.1 |

**Table S2.**
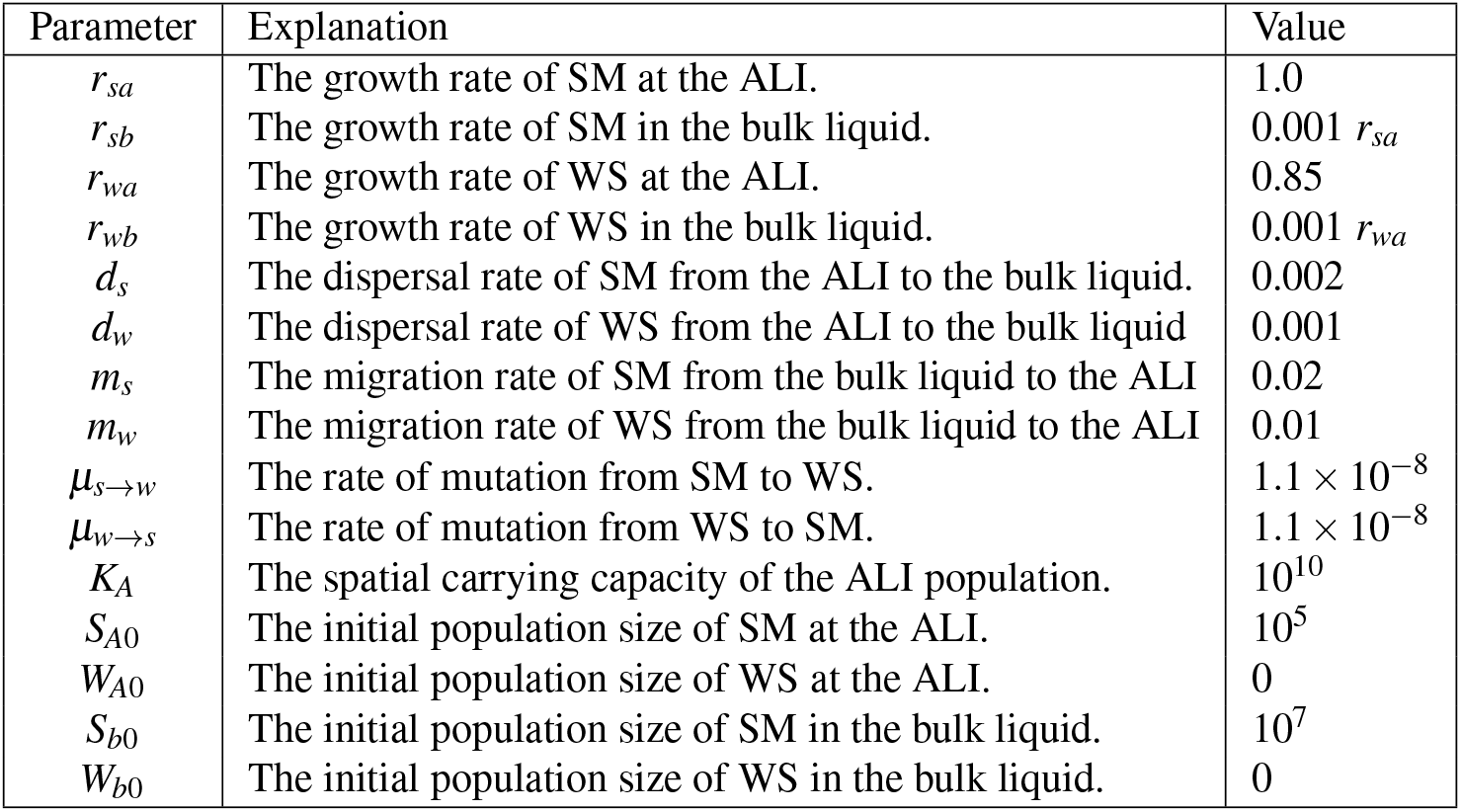
List of parameters in the four-state model simulations.

| Parameter | Explanation | Value |
| --- | --- | --- |
| $r_{sa}$ | The growth rate of SM at the ALI. | 1.0 |
| $r_{sb}$ | The growth rate of SM in the bulk liquid. | $0.001 r_{sa}$ |
| $r_{wa}$ | The growth rate of WS at the ALI. | 0.85 |
| $r_{wb}$ | The growth rate of WS in the bulk liquid. | $0.001 r_{wa}$ |
| $d_s$ | The dispersal rate of SM from the ALI to the bulk liquid. | 0.002 |
| $d_w$ | The dispersal rate of WS from the ALI to the bulk liquid | 0.001 |
| $m_s$ | The migration rate of SM from the bulk liquid to the ALI | 0.02 |
| $m_w$ | The migration rate of WS from the bulk liquid to the ALI | 0.01 |
| $\mu_{s \rightarrow w}$ | The rate of mutation from SM to WS. | $1.1 \times 10^{-8}$ |
| $\mu_{w \rightarrow s}$ | The rate of mutation from WS to SM. | $1.1 \times 10^{-8}$ |
| $K_A$ | The spatial carrying capacity of the ALI population. | $10^{10}$ |
| $S_{A0}$ | The initial population size of SM at the ALI. | $10^5$ |
| $W_{A0}$ | The initial population size of WS at the ALI. | 0 |
| $S_{b0}$ | The initial population size of SM in the bulk liquid. | $10^7$ |
| $W_{b0}$ | The initial population size of WS in the bulk liquid. | 0 |

**Table S3.** The list of *Pseudomonas fluorescens* SBW25 strains used in this paper. The mutant strains without references were obtained from the Microbial Population Biology strain repository (Max Planck Institute for Evolutionary Biology, Germany), and all were constructed by a two-step allelic exchange protocol. *wspF, awsX*, and *mwsR* are negative regulators of c-i-GMP^37^, and mutations on the loci typically lead to the overexpression of c-di-GMP. Deleting *wss* disables the production of cellulose, the main extracellular matrix of *P. fluorescens* SBW25. Deleting *fliA* makes cells immotile, and deleting *pvds* eliminates the expression of the fluorescent siderophore pyoverdin. ^∗^ A mat is successfully formed only from a very high inoculum (∼10^8^ cells/mL). ^∗∗^ mScarlet is integrated from pMRE-Tn7-145 (Addgene 118561), and sGFP2 is integrated from pMRE-Tn7-152 (Addgene 118566)^53^. ^∗∗∗^ mScarlet and sGFP2 are integrated in Δ*pvdS* strains from^54^.

| Strain | Mat | Genotype | Internal ID | Reference |
| --- | --- | --- | --- | --- |
| SBW25 wild type (SM) | No |  | MPB13805 (PBR341) | 1,50,51 |
| Wrinkly spreader (WS) *wspF* | Yes | *wspF* (PFLU1224) Δ150bp (677-826). | MPB08384 | 41 |
| Wrinkly spreader (WS) *awsX* | Yes | *awsX* (PFLU5211) Δ32bp (229-261). | MPB10583 | 37 |
| Wrinkly spreader (WS) *mwsR* | Yes | *mwsR* (PFLU5329) T2183C. | MPB10613 | 37 |
| WS(*wspF*)-Δ*fliA* | Yes* | *wspF* (PFLU1224) Δ150bp (677-826), Δ*fliA* (PFLU4417). | MPB36210 | This work. |
| SM-Δ*fliA* | No | Δ*fliA* (PFLU4417). | MPB14427 | This work. |
| SM-Δ*wss* | No | Δ*wss* (PFLU0300-0309). | MPB14424 | This work. |
| WS(*wspF*)-mScarlet | Yes | *wspF* (PFLU1224) Δ150bp (677-826), attTn7::Tn7-PnptII-mScarlet-I-Gm. | MPB16290 | This work**. |
| WS(*wspF*)-Δ*pvdS*-sGFP2 | Yes | *wspF* (PFLU1224) Δ150bp (677-826), Δ*pvdS* (PFLU4389), attTn7::Tn7-PnptII-sGFP2-Kan. | MPB23958 | This work***. |
| WS(*wspF*)-Δ*pvdS*-mScarlet | Yes | *wspF* (PFLU1224) Δ150bp (677-826), Δ*pvdS* (PFLU4389), attTn7::Tn7-PnptII-mScarlet-I-Gm. | MPB23959 | This work***. |
| SM-mScarlet | No | attTn7::Tn7-PnptII-mScarlet-I-Gm. | MPB15845 | 52 |
| SM-Δ*pvdS*-sGFP2 | No | Δ*pvdS* (PFLU4389), attTn7::Tn7-PnptII-sGFP2-Kan. | MPB26812 | This work*** |
| SM-Δ*pvdS*-mScarlet | No | Δ*pvdS* (PFLU4389), attTn7::Tn7-PnptII-mScarlet-I-Gm. | MPB26813 | This work***. |
| SM pCdrA-mScarlet | No | pCdrA-mScarlet (modified from Addgene 111614): plasmid cdrA-RBSII-mScarlet-T0-T1, pRO1600, oriV; GmR and AmR<sup>25</sup> | MPB38082 | This work. |

## Supplementary figures

**Figure S1.**
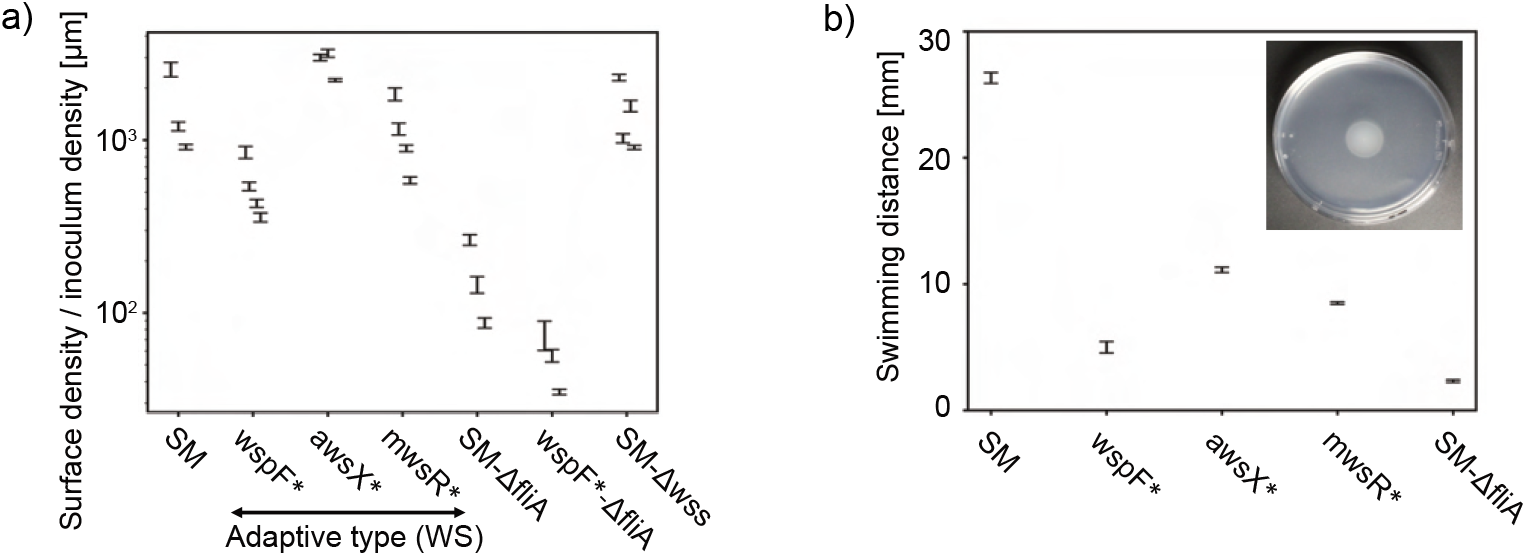
Measurements of motility and early ALI occupation with various strains. (a) The efficiency of ALI occupation was quantified by the ratio of the surface density at *t* =2 h over the inoculum volume density. The surface density was measured by counting the number of cells at the ALI with microscopy. 3 or 4 inoculum densities were tested for each strain, shown as different points in the plot (from left to right, the inoculum densities are low to high). (b) Motility assay on semi-solid agar plates. The inset shows the representative motility assay on a 10 cm semi-solid agar plate.

**Figure S2.**
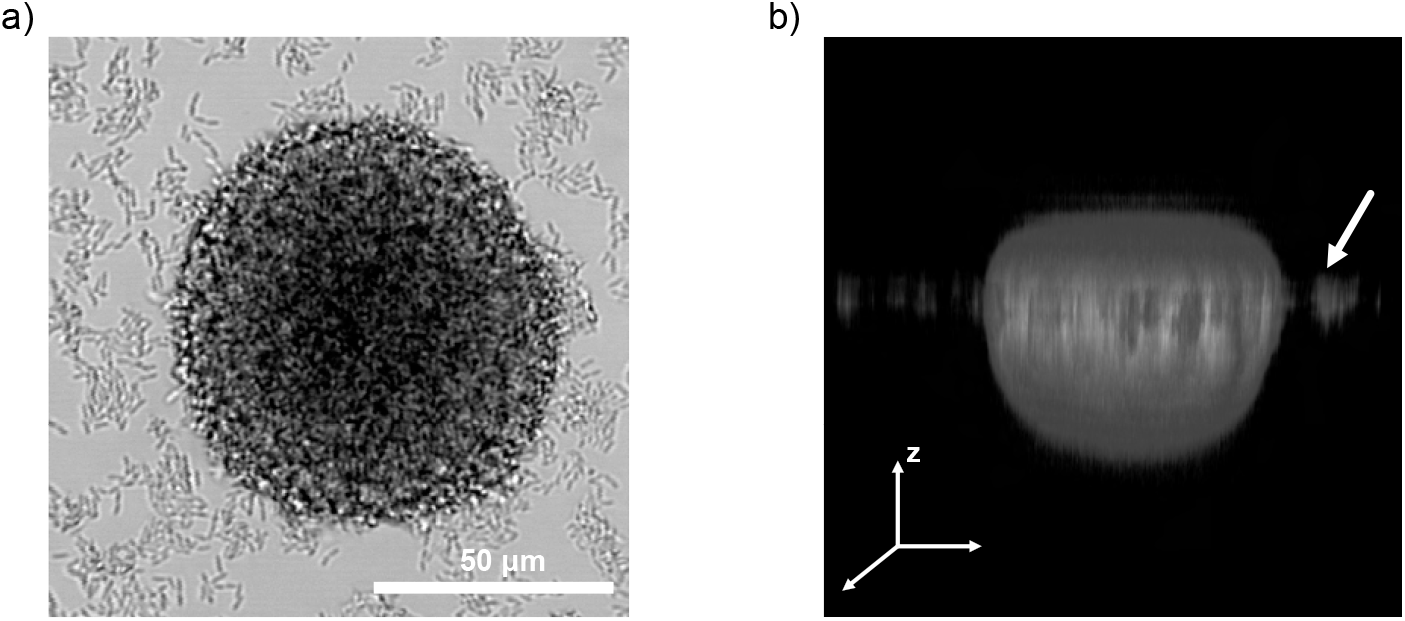
3D structure of a WS microcolony at an ALI. (a) A top view of a microcolony at an air-liquid interface. (b) The 3D reconstruction of the microcolony in (a) by a confocal microscope. The z-position of small cell clusters (white arrow) was lower than the top surface of the microcolony, suggesting that the microcolony penetrated the air-liquid interface.

**Figure S3.**
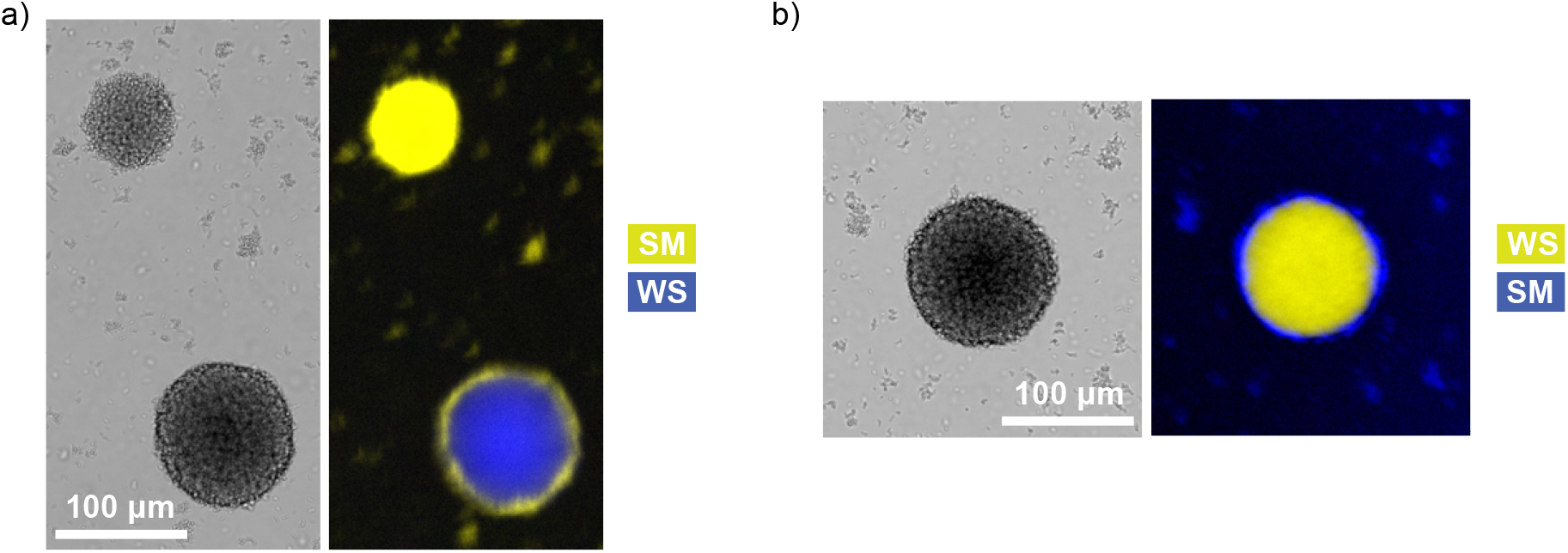
Adhesion of the wild type strain (SM) to microcolonies of the mat-forming strain (WS). The SM and WS strains were co-cultured starting from a low inoculum (∼5 ×10^3^ cells/mL for each), and imaged by fluorescent microscopy at *t* =16 h. The flipped combinations of the fluorescence proteins were shown in (a) (SM-mScarlet, WS-sGFP2) and (b) (SM-sGFP2, WS-mScarlet).

**Figure S4.**
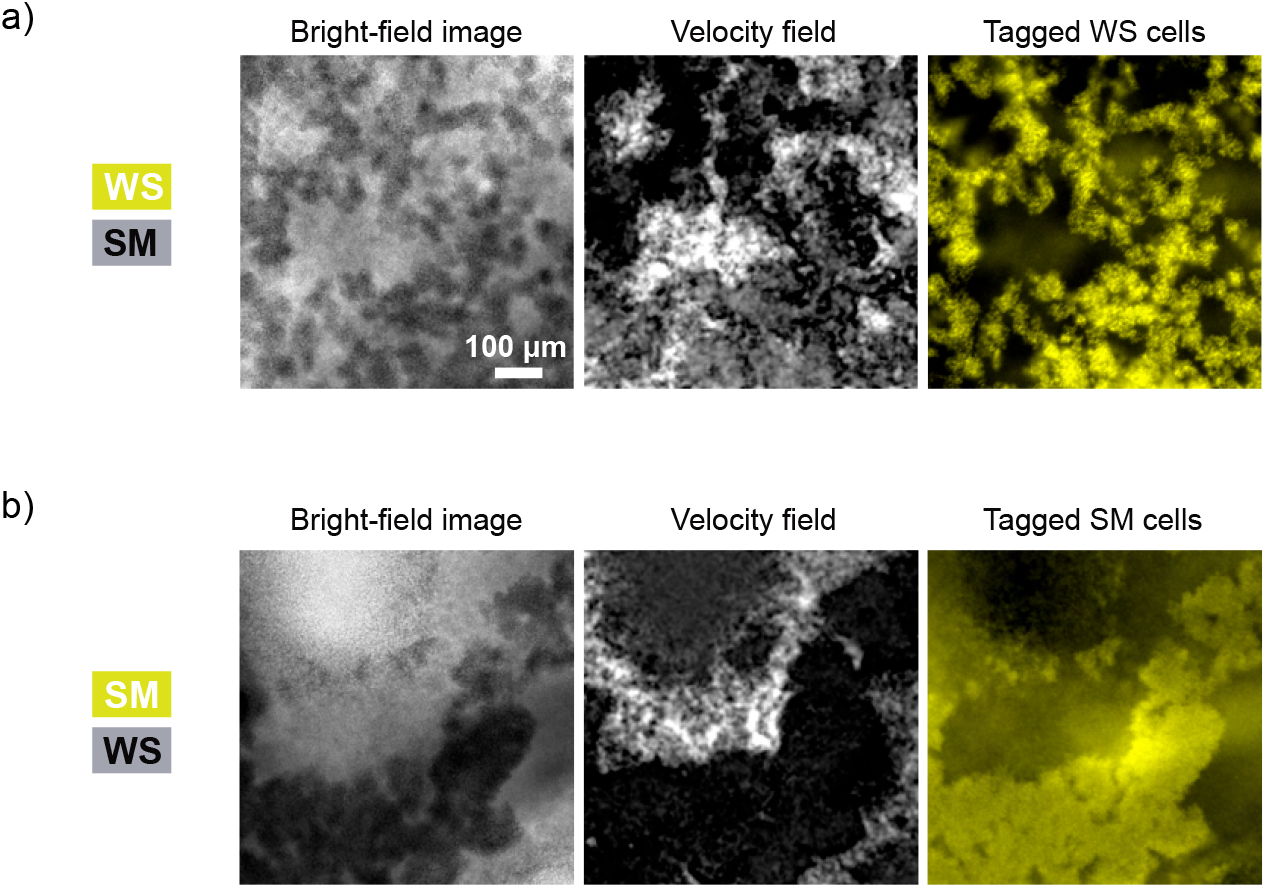
Inclusion of SM in the sessile regions during transition to motility at the ALI. Snapshots and PIV velocity fields were shown for co-cultures of SM and WS-mScarlet (a) and SM-mScarlet and WS (b). For the velocity field, the absolute value of the velocity at each position was averaged over 5 seconds. The images were taken at *t* =10.5 h after inoculation.

## Supplementary Movie Captions

### Supplementary Movie 1

Inoculum-dependent failure of mat formation by an adaptive Wrinkly Spreader type (WS). Mono-culture of WS could not form a mat structure when started from a low inoculum density (dilution factor 10^4^). 3 mL of the cell suspension was statically cultured in a well of a 12-well plate at 28 ^*°*^C. The movie was taken with a stereo microscope (Axio Zoom V16, Zeiss) in dark-field illumination with 16X magnification. The time unit is [hour:minute].

### Supplementary Movie 2

Successful mat formation by WS. Mono-culture of WS could develop a robust mat when started from a high inoculum density (dilution factor 10^2^). 3 mL of the cell suspension was statically cultured in a well of a 12-well plate at 28 ^*°*^C. The movie was taken with a stereo microscope (Axio Zoom V16, Zeiss) in dark-field illumination with 16X magnification. The time unit is [hour:minute].

### Supplementary Movie 3

Representative observation of microcolony collapse at the air-liquid interface with high magnification. WS microcolony collapse was captured at the center of a well with 140X magnification. 3 mL of WS low-inoculum culture (dilution factor 10^7^) was statically cultured in a well of a 12-well plate at 28 ^*°*^C. The movie was taken with a stereo microscope (Axio Zoom V16, Zeiss) in dark-field illumination.

### Supplementary Movie 4

Phase separation of active fluid-like regions and sessile regions at the air-liquid interface of a co-culture of SM and WS. A mixture of SM and WS was cultured under static conditions. The movie was taken at *t* =10.5 h for 5 seconds with bright-field imaging. The same data is also shown in Fig. S4a.

## Notes

https://doi.org/10.5281/zenodo.15293177

https://doi.org/10.6019/S-BIAD2117

### Competing Interest Statement

The authors have declared no competing interest.

### Summary of Updates

Strain list (Table S3) was extended to include comprehensive information; References were extended; Figures were modified to improve the clarity; Typos were corrected; Data availability statement was updated to include the link to scripts, spreadsheets, and microscopy data; Author list were updated to include ORCID.

## References

1. Rainey, P. B. & Travisano, M. Adaptive radiation in a heterogeneous environment. Nature 394, 69–72 (1998).

2. Kovács, Á. T. & Dragoš, A. Evolved biofilm: review on the experimental evolution studies of bacillus subtilis pellicles. J. molecular biology 431, 4749–4759 (2019).

3. Pentz, J. T. & Lind, P. A. Forecasting of phenotypic and genetic outcomes of experimental evolution in pseudomonas protegens. PLoS Genet. 17, e1009722 (2021).

4. Smith, T. M. et al. Rapid adaptation of a complex trait during experimental evolution of mycobacterium tuberculosis. Elife 11, e78454 (2022).

5. Pentz, J. T., Biswas, A., Alsaed, B. & Lind, P. A. Forecasting of phenotypic and genetic outcomes of experimental evolution in pseudomonas syringae and pseudomonas savastanoi. bioRxiv 2024–02 (2024).

6. Koza, A., Moshynets, O., Otten, W. & Spiers, A. J. Environmental modification and niche construction: developing o2 gradients drive the evolution of the wrinkly spreader. The ISME journal 5, 665–673 (2011).

7. Hengge, R. Principles of c-di-gmp signalling in bacteria. Nat. Rev. Microbiol. 7, 263–273 (2009).

8. Guttenplan, S. B. & Kearns, D. B. Regulation of flagellar motility during biofilm formation. FEMS microbiology reviews 37, 849–871 (2013).

9. Valentini, M. & Filloux, A. Biofilms and cyclic di-gmp (c-di-gmp) signaling: lessons from pseudomonas aeruginosa and other bacteria. J. Biol. Chem. 291, 12547–12555 (2016).

10. Lind, P. A., Libby, E., Herzog, J. & Rainey, P. B. Predicting mutational routes to new adaptive phenotypes. Elife 8, e38822 (2019).

11. Rainey, P. B. & Rainey, K. Evolution of cooperation and conflict in experimental bacterial populations. Nature 425, 72–74 (2003).

12. Jerdan, R., Kuśmierska, A., Petric, M. & Spiers, A. J. Penetrating the air–liquid interface is the key to colonization and wrinkly spreader fitness. Microbiology 165, 1061–1074 (2019).

13. Ardré, M., Dufour, D. & Rainey, P. B. Causes and biophysical consequences of cellulose production by pseudomonas fluorescens sbw25 at the air-liquid interface. J. bacteriology 201, 10–1128 (2019).

14. Hammerschmidt, K., Rose, C. J., Kerr, B. & Rainey, P. B. Life cycles, fitness decoupling and the evolution of multicellularity. Nature 515, 75–79 (2014).

15. Rainey, P. B. & Kerr, B. Cheats as first propagules: a new hypothesis for the evolution of individuality during the transition from single cells to multicellularity. BioEssays 32, 872–880 (2010).

16. Giddens, S. R. et al. Mutational activation of niche-specific genes provides insight into regulatory networks and bacterial function in a complex environment. Proc. Natl. Acad. Sci. 104, 18247–18252 (2007).

17. Spiers, A. J. & Rainey, P. B. The pseudomonas fluorescens sbw25 wrinkly spreader biofilm requires attachment factor, cellulose fibre and lps interactions to maintain strength and integrity. Microbiology 151, 2829–2839 (2005).

18. Hölscher, T. et al. Motility, chemotaxis and aerotaxis contribute to competitiveness during bacterial pellicle biofilm development. J. molecular biology 427, 3695–3708 (2015).

19. Qin, B. et al. Hierarchical transitions and fractal wrinkling drive bacterial pellicle morphogenesis. Proc. Natl. Acad. Sci. 118, e2023504118 (2021).

20. Loferer-Krossbacher, M., Klima, J. & Psenner, R. Determination of bacterial cell dry mass by transmission electron microscopy and densitometric image analysis. Appl. environmental microbiology 64, 688–694 (1998).

21. Milo, R. & Phillips, R. Cell biology by the numbers (Garland Science, 2015).

22. Chen, W., Lickfield, G. C. & Yang, C. Q. Molecular modeling of cellulose in amorphous state. part i: model building and plastic deformation study. Polymer 45, 1063–1071 (2004).

23. Mazeau, K. & Heux, L. Molecular dynamics simulations of bulk native crystalline and amorphous structures of cellulose. The J. Phys. Chem. B 107, 2394–2403 (2003).

24. Davis, S., Trapman, P., Leirs, H., Begon, M. & Heesterbeek, J. The abundance threshold for plague as a critical percolation phenomenon. Nature 454, 634–637 (2008).

25. Rybtke, M. T. et al. Fluorescence-based reporter for gauging cyclic di-gmp levels in pseudomonas aeruginosa. Appl. environmental microbiology 78, 5060–5069 (2012).

26. Summers, J. Evolution of developmental regulation in a simple multicellular life cycle. Ph.D. thesis, Christian-Albrechts-Universität Kiel; Plön (2023).

27. Gehrig, S. Adaptation of Pseudomonas fluorescens SBW25 to the air-liquid interface: a study in evolutionary genetics. Ph.D. thesis, University of Oxford (2005).

28. Sauer, K. et al. The biofilm life cycle: expanding the conceptual model of biofilm formation. Nat. Rev. Microbiol. 20, 608–620 (2022).

29. Fukami, T., Beaumont, H. J., Zhang, X.-X. & Rainey, P. B. Immigration history controls diversification in experimental adaptive radiation. Nature 446, 436–439 (2007).

30. Theodosiou, L., Farr, A. D. & Rainey, P. B. Barcoding populations of pseudomonas fluorescens sbw25. J. Mol. Evol. 91, 254–262 (2023).

31. Spiers, A. J., Kahn, S. G., Bohannon, J., Travisano, M. & Rainey, P. B. Adaptive divergence in experimental populations of pseudomonas fluorescens. i. genetic and phenotypic bases of wrinkly spreader fitness. Genetics 161, 33–46 (2002).

32. Bittleston, L. S., Gralka, M., Leventhal, G. E., Mizrahi, I. & Cordero, O. X. Context-dependent dynamics lead to the assembly of functionally distinct microbial communities. Nat. communications 11, 1440 (2020).

33. Boyer, S., Hérissant, L. & Sherlock, G. Adaptation is influenced by the complexity of environmental change during evolution in a dynamic environment. PLoS Genet. 17, e1009314 (2021).

34. Karve, S. & Wagner, A. Environmental complexity is more important than mutation in driving the evolution of latent novel traits in e. coli. Nat. communications 13, 5904 (2022).

35. Rosenzweig, R. F., Sharp, R., Treves, D. S. & Adams, J. Microbial evolution in a simple unstructured environment: genetic differentiation in escherichia coli. Genetics 137, 903–917 (1994).

36. Treves, D. S., Manning, S. & Adams, J. Repeated evolution of an acetate-crossfeeding polymorphism in long-term populations of escherichia coli. Mol. biology evolution 15, 789–797 (1998).

37. McDonald, M. J., Gehrig, S. M., Meintjes, P. L., Zhang, X.-X. & Rainey, P. B. Adaptive divergence in experimental populations of pseudomonas fluorescens. iv. genetic constraints guide evolutionary trajectories in a parallel adaptive radiation. Genetics 183, 1041–1053 (2009).

38. Lind, P. A., Farr, A. D. & Rainey, P. B. Experimental evolution reveals hidden diversity in evolutionary pathways. elife 4, e07074 (2015).

39. Lind, P. A., Farr, A. D. & Rainey, P. B. Evolutionary convergence in experimental pseudomonas populations. The ISME J. 11, 589–600 (2017).

40. King, E. O., Ward, M. K. & Raney, D. E. Two simple media for the demonstration of pyocyanin and fluorescin. The J. laboratory clinical medicine 44, 301–307 (1954).

41. Bantinaki, E. et al. Adaptive divergence in experimental populations of pseudomonas fluorescens. iii. mutational origins of wrinkly spreader diversity. Genetics 176, 441–453 (2007).

42. Thielicke, W. & Sonntag, R. Particle image velocimetry for matlab: Accuracy and enhanced algorithms in pivlab. J. Open Res. Softw. 9, 12 (2021).

43. Eden, M. A two-dimensional growth process. Dyn. fractal surfaces 4, 223–239 (1961).

44. Gibson, M. A. & Bruck, J. Efficient exact stochastic simulation of chemical systems with many species and many channels. The journal physical chemistry A 104, 1876–1889 (2000).

45. Anderson, D. F. A modified next reaction method for simulating chemical systems with time dependent propensities and delays. The J. chemical physics 127 (2007).

46. Atis, S., Weinstein, B. T., Murray, A. W. & Nelson, D. R. Microbial range expansions on liquid substrates. Phys. review X 9, 021058 (2019).

47. Pirt, S. A kinetic study of the mode of growth of surface colonies of bacteria and fungi. Microbiology 47, 181–197 (1967).

48. Grimson, M. J. & Barker, G. C. Continuum model for the spatiotemporal growth of bacterial colonies. Phys. Rev. E 49, 1680 (1994).

49. Cordero, M., Mitarai, N. & Jauffred, L. Motility mediates satellite formation in confined biofilms. The ISME J. 1–9 (2023).

50. Rainey, P. B. & Bailey, M. J. Physical and genetic map of the pseudomonas fluorescens sbw25 chromosome. Mol. microbiology 19, 521–533 (1996).

51. Silby, M. W. et al. Genomic and genetic analyses of diversity and plant interactions of pseudomonas fluorescens. Genome biology 10, 1–16 (2009).

52. Yulo, P. R. J. et al. Evolutionary rescue of spherical mreb deletion mutants of the rod-shape bacterium pseudomonas fluorescens sbw25. eLife 13, RP98218 (2025).

53. Schlechter, R. O. et al. Chromatic bacteria–a broad host-range plasmid and chromosomal insertion toolbox for fluorescent protein expression in bacteria. Front. Microbiol. 9, 3052 (2018).

54. Moreno-Fenoll, C., Ardré, M. & Rainey, P. B. Polar accumulation of pyoverdin and exit from stationary phase. Microlife 5, uqae001 (2024).

